# The Effect of Gelling agent, medium pH and silver nitrate on adventitious shoot regeneration in *Solanum tuberosum*

**DOI:** 10.1101/2020.01.03.894063

**Authors:** Amanpreet Kaur, Anil Kumar

## Abstract

Vegetative propagation of potato makes the crop vulnerable to many seed borne diseases. The importance of the crop in attainment of food security makes it an important candidate for in vitro propagation and genetic manipulations. To undertake crop improvement programmes, development of an efficient regeneration protocol is a pre-requisite. Therefore, the present report was focussed to study various factors affecting shoot organogenesis in potato cultivar ‘Kufri Chipsona 1’. The incorporation of silver nitrate (10 µM) to the regeneration medium (MS medium supplemented with BA and GA_3_) was found to induce shoot organogenesis in 32.11% of leaf and 59.99% of internodal explants. An increase in mean number of shoots regenerated per leaf (5.31) and internodal (8.67) explant was also observed upon addition of silver nitrate to the medium. Similarly, solidification of medium with clarigel and its adjustment to pH 5.8 was found optimum for increasing shoot organogenesis frequency in potato. Among the two types of explants tested, a better response was observed from internodes in comparison with leaf explants. The regenerated shoots were tested for clonal fidelity using PCR based molecular markers [Random Amplified Polymorphic DNA (RAPD) and Inter-Simple Sequence Repeats (ISSR)] and were found true to type.

## Introduction

Organogenesis through adventitious shoot regeneration is an important phenomena of plant tissue culture which acts as a first step in cultivar improvement through genetic transformation (Tsao and Reed 2002). However, the efficiency of shoot organogenesis is largely dependent on factors such as culture conditions and medium composition (Kumar 1996, Kumar et al. 2003). In potato, an important rabi food crop, shoot organogenesis is extensively studied using different explants such as leaf, internode and tuber discs (Yasmin et al. 2003, Elaleem et al. 2009, Molla et al. 2011, Rezende et al. 2013, Ghosh et al. 2014, Abdelaleem 2015, Campos et al. 2016, Kaur et al. 2017). Most of the reports are mainly focussed on Plant Growth regulators (PGRs), but, studies on the role of other factors such as gelling agents, light intensity, medium pH, silver nitrate etc. in shoot organogenesis remains very limited (Cui et al. 2000, Kumar et al. 2003, Kumar et al. 2010, Yaacob et al. 2014). Medium pH and gelling agents are seen as one of the most important factors due to their role in regulation of medium nutrient solubility and their uptake by the explants (George 1993, Bhatia and Ashwath 2005). In real terms, the effect of medium pH on cytoplasm of a cell is not long lasting as cells are able to readjust external pH by preferential uptake of ions during changed environmental conditions (uptake of anions or cations is increased if medium pH is low or high respectively) (George 1993, Parton et al. 1997). Thus the explant death occurring as a result of altered medium pH is not due to cell damage but due to nutrient imbalances. At low pH, inorganic phosphates get converted into organic phosphates, thus, reducing Adenosine triphosphate (ATP) levels in extracellular regions of the cell (Mimura et al. 2000). pH also plays an important role is solidification of the medium. It has been reported that pH higher than 6.0 results in more solidification of the medium which makes nutrient availability scarce for the plant (Owen et al. 1991). In general, Agar is used as a most common gelling agent but problems such as variability among batches, vitrification, presence of impurities and growth inhibitory compounds limits its use in propagation medium (Stolz 1971, Debergh et al. 1992, Nairn et al. 1995). These limitations may be overcome by other gelling agents such as gellan gums (gelrite and phytagel) having high ash content, low impurities and more consistancy (Huang et al. 1995). Therefore, the choice of gelling agent for efficient regeneration system development is an important aspect of the study.

In plant tissue culture, the use of sealed propagation vessels lead to accumulation of ethylene upto active physiological levels (Righetti et al. 1990, Matthys et al. 1995). This can induce or inhibit differentiation of plant cells into adventitious shoots (Gonzalez et al. 1997). The use of ethylene biosynthesis or action inhibitors such as silver nitrate, silver thiosulphate, aminoethoxyvinylglycine have shown improvement in shoot organogenic efficiencies in many plants including potato (Naik and Chand 2003, Mookkan and Andy 2014, Kaur et al. 2017). However, it is very important to note that the concentration of the inhibitor in the medium plays an important role in morphogenesis by regulating ethylene levels optimum for shoot differentiation.

The present study independently assesses the effect of medium pH (5.2-7.0), gelling agent (gelrite, agar, clarigel) and silver nitrate (0-20µM) on shoot regeneration of potato cultivar ‘Kufri Chipsona 1’. The effect of above mentioned factors on variation in chlorophyll content was also studied.

## Material and Methods

### Plant material and medium composition

The microshoot cultures of *Solanum tuberosum* L. cultivar ‘Kufri Chipsona 1’ (CS-1) maintained at Thapar Institute of Engineering & Technology, Patiala, were subcultured using nodal explants at the regular interval of 21 days on MS1 medium (Kaur et al. 2017; Murashige and Skoog (1962) medium supplemented with 10 µM silver nitrate (AgNO_3_), 74mM sucrose and 0.7% (w/v) agar as gelling agent) to obtain fully expanded leaves for experimentation. All the experiments were carried out in glass culture bottles (Kasablanka, Mumbai) containing 30ml regeneration medium (Kaur et al. 2017; MS1 medium supplemented 10 µM BA and 15 µM GA_3_), here onwards referred to as MS2 medium. All tissue culture grade chemicals were purchased from HiMedia Laboratories, Mumbai, India.

### Effects of Silver Nitrate (AgNO_3_), gelling agents and pH on shoot organogenesis

The effect of AgNO_3_, gelling agents and medium pH were evaluated independently. Leaf and internodal segments excised from 3-4 weeks old microshoots of CS-1 were cultured horizontally (for internodes) or with adaxial side in contact (for leaf explants) with MS2 solidified either with Agar-agar (0.7% w/v), Clarigel (0.25% w/v), Gelrite (0.35%w/v) and adjusted to pH 5.2, 5.5, 5.8, 6.1, 6.4, 6.7 and 7.0 to evaluate the effect of gelling agent and medium pH on organogenic potential of the potato explants.

To investigate the role of AgNO_3_ in shoot organogenesis, leaf and internodal explants of CS-1 were inoculated on MS medium containing 10 µM BA and 15 µM GA_3_ and supplemented with different concentrations of AgNO_3_ (0, 5, 10, 15, 20 µM). AgNO_3_ was added to the medium before autoclaving. Cultures were maintained at 25±2°C under 16/8 h light/dark conditions at photon flux intensity of 50 µmol m^-2^s^-1^ provided by cool white fluorescent lamps (Philips India, Mumbai). Data were recorded after 4-6 weeks.

### Clonal fidelity studies

Clonal studies were carried out using random amplified polymorphic DNA (RAPD) and inter-simple sequence repeat (ISSR) markers. Regenerated shoots (randomly selected) were multiplied on MS1 medium and total genomic DNA was isolated using method described by Doyle and Doyle (1990). PCR amplifications were carried out using five each of RAPD and ISSR primers using previously described amplification conditions (Kaur et al. 2017). Amplified products were separated on ethidium bromide stained agarose gel 1.2% (w/v) and visualised under UV trans-illuminator (Gel Doc Mega; Biosystematica, USA).

### Estimation of Chlorophyll content

Chlorophyll content (Chlorophyll a, b and total content) was estimated according to the method described by Arnon (1949). Plant tissue sample (200 mg) was collected after four weeks of culture on various media combinations and were homogenised in 80% (v/v) aqueous solution of acetone. After centrifugation at 10000g for 20 minutes, pellet was re-extracted till it becomes colourless. Supernatants were pooled to make total volume upto 20ml. Absorbance was measured at 663nm and 645nm spectrophotometrically. Chlorophyll a and b (mg/l) was measured using formula (12.7_A663_-2.7_A645_) and (22.9_A645_-4.7_A663_) respectively.

### Statistical analysis

Each experiment consisted of five replicates with five explants in each culture vessel and was repeated twice. Chlorophyll content was estimated three times for same sample and was repeated thrice. Data were analysed through one way Analysis of variance (ANOVA). Significant differences between means were compared using Duncan’s multiple range test at 0.05 value of P using CoStat (CoHort, USA). Graphical analysis was carried out using Graph Pad Prism 5 software (Graph Pad, CA).

## Results

Gelling agent (agar, gelrite, clarigel), incorporation of silver nitrate in regeneration medium and pH were found to have an important role in induction of shoot organogenesis in potato (Figure 1A-B). After four weeks of culture, green coloured callus was induced from mid-rib region of leaf explants and cut ends of internodal explants in all the media combinations and shoots were regenerated from this callus after six weeks.

**Figure 1:**
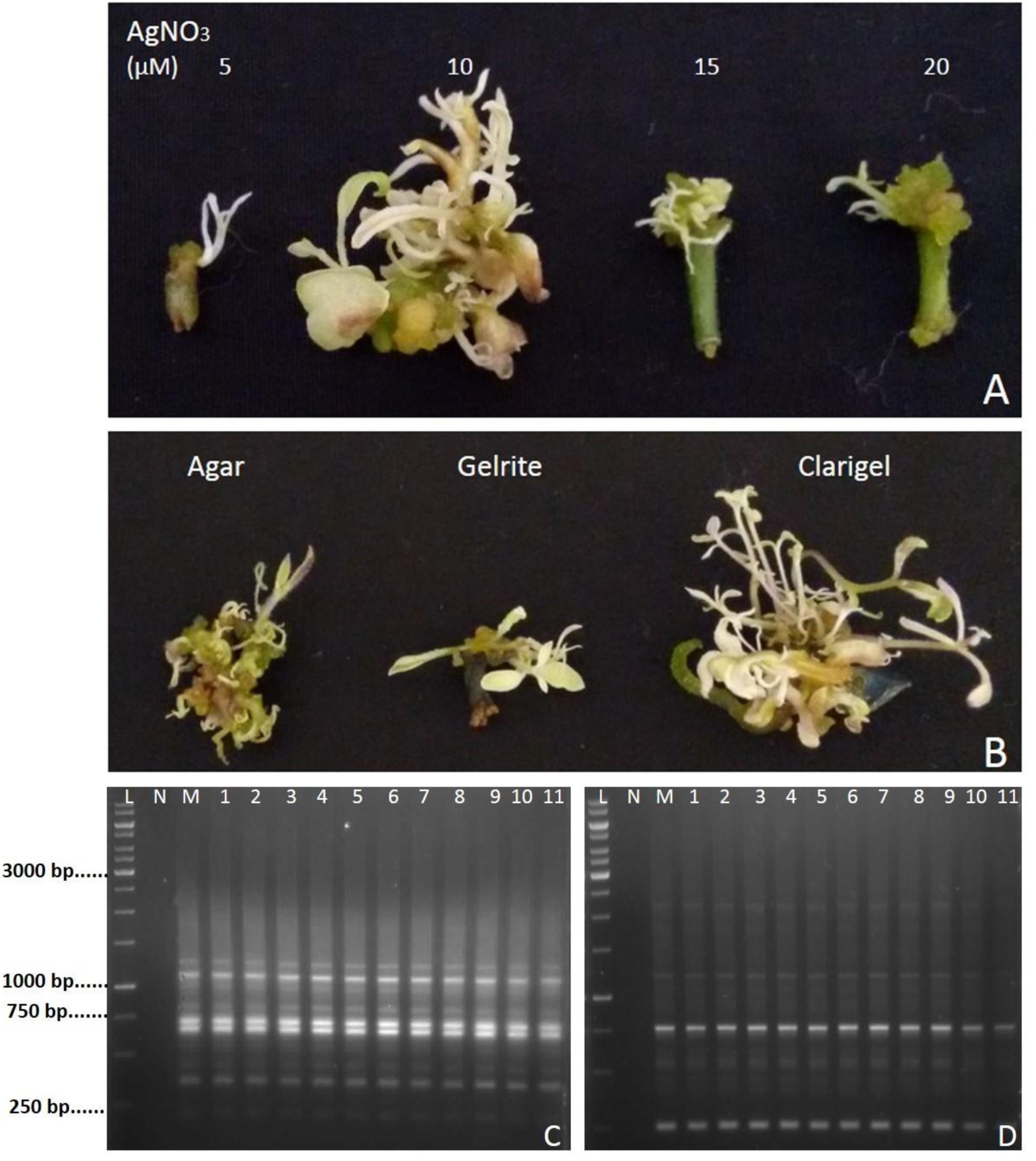
Shoot organogenesis and clonal fidelity studies in *Solanum tuberosum* cultivar ‘Kufri Chipsona 1’. **A-B;** Variation in shoot induction from callus induced on medium supplemented with BA (10 µM), GA_3_ (15 µM), **A;** containing different concentrations of silver nitrate and **B;** solidified with various gelling agents (Agar-Agar, clarigel and gelrite) **C-D;** RAPD and ISSR profiles of DNA isolated from regenerated shoots using primers **C;** RAPD 4 and **D;** ISSR20. *Lanes: M* mother culture; *1-11* lines of regenerated shoots; *L* Marker, *N* Negative control.

### Effect of gelling agent

Gelling agent was found to significantly affect organogenesis in potato cultivar CS-1 (Figure 1 and 2). Maximum shoot regeneration was observed from leaf (41.07%) and internodal (49.96%) explants on MS2 medium solidified with clarigel (Figure 2A). But maximum number of shoots regenerated per explant was recorded on medium solidified with agar (Figure 2B).

**Figure 2:**
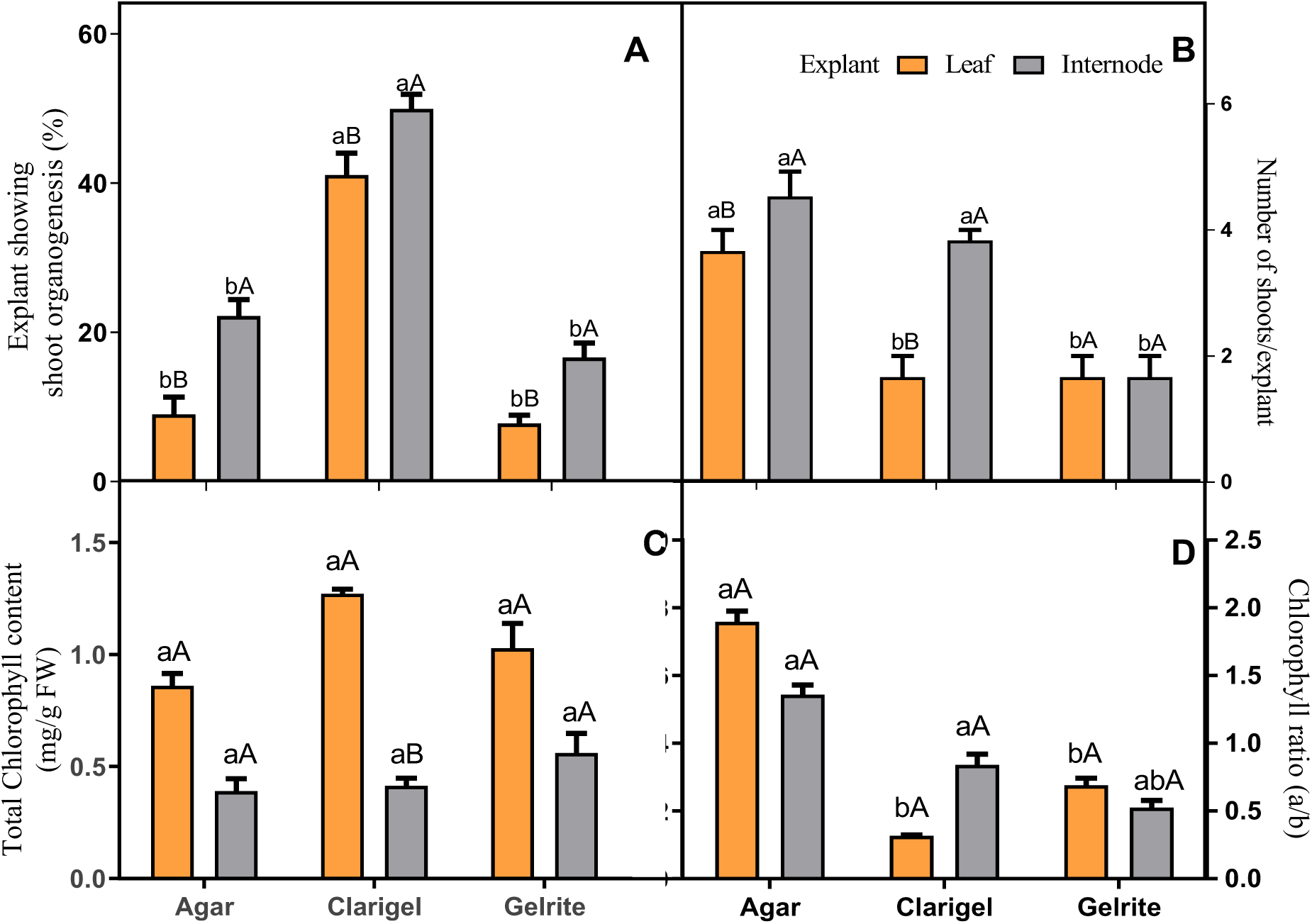
The effect of gelling agent (Agar, clarigel, gelrite) on organogenic efficiencies and associated chlorophyll changes in leaf and internodal explants taken from potato cultivar ‘Kufri Chipsona 1’. Data were recorded after 6 weeks and analysed using ANOVA. Mean values within the explants were compared using Duncan’s multiple range test (DMRT) and between the explants were compared using unpaired t-test at P<0.05. Bars with same lowercase letter (within explants) and uppercase letter (between explants) was non-significant at P<0.05.

Total chlorophyll content was also found to vary between the explants cultured on MS2 medium solidified with different gelling agents (Figure 2C). However the variation was not significant except for explants cultured on medium solidified with clarigel. Highest chlorophyll a/b ratio was found on medium solidified with agar for both the explants (Figure 2D).

### Effect of medium pH

The pH of medium was found to drastically influence the shoot organogenesis in leaf and internodal explants of CS-1. Both explants showed maximum organogenic response on MS2 medium with pH adjusted in the range of 5.3-5.8. Any further change in pH led to decrease in organogenic potential of both explants (Figure 3). Leaf explants were found to be more sensitive towards pH change as almost negligible shoot organogenesis was observed at pH above 6.3. In contrast, about 22.23% internodal explants showed shoot organogenesis even on MS2 medium with pH value of 7.3 (Figure 3A).

**Figure 3:**
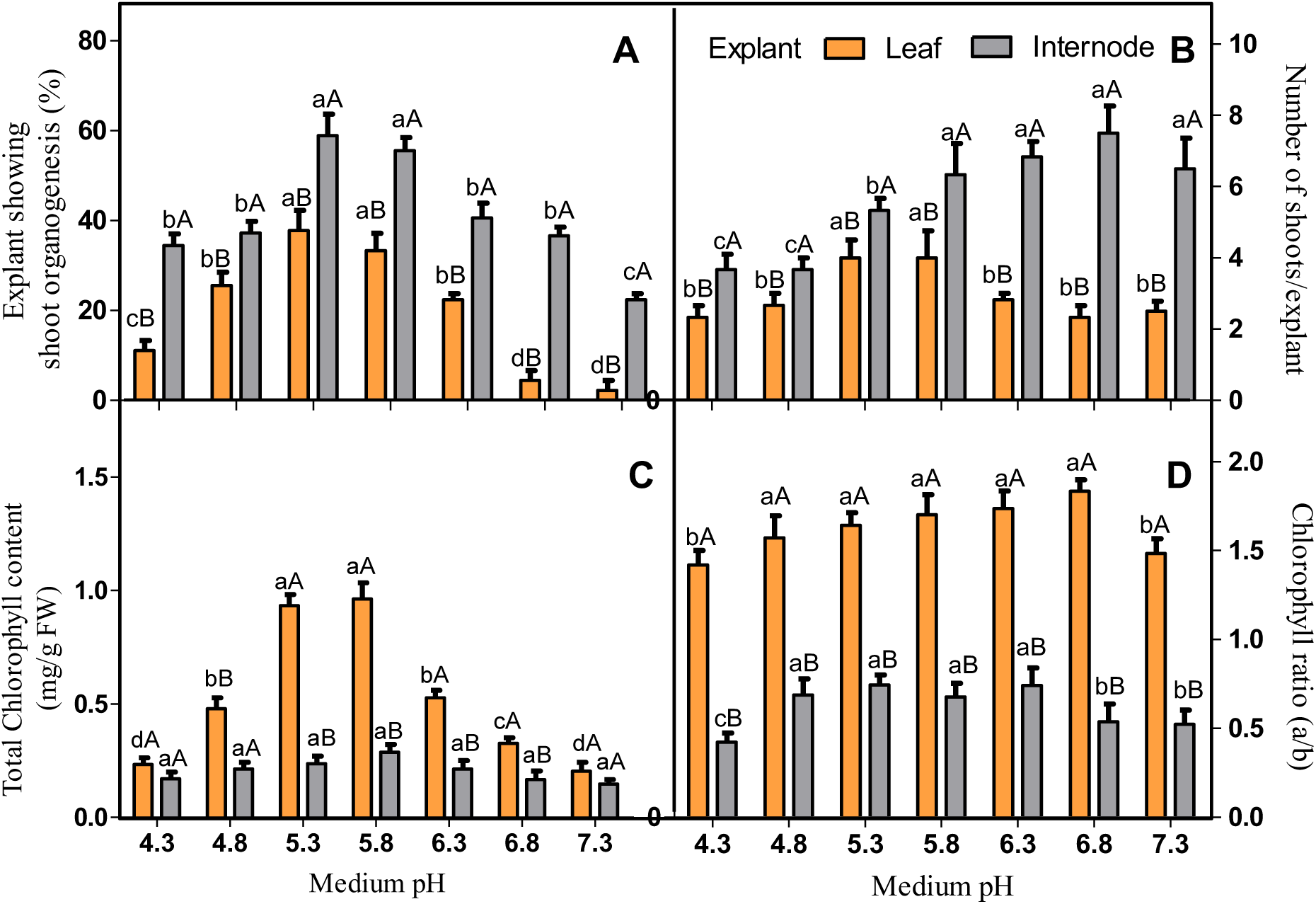
The effect of pH on organogenic efficiencies and associated chlorophyll changes in leaf and internodal explants taken from potato cultivar ‘Kufri Chipsona 1’. Data were recorded after 6 weeks and analysed by ANOVA. Mean values within the explant and between the explants were compared using Duncan’s multiple range test (DMRT) and unpaired t-test respectively. Bars with same lowercase letter (within explants) and uppercase letter (between explants) was non-significant at P<0.05.

Mean number of shoots regenerated per explant were also found to vary with change in medium pH (Figure 3B). Increase in pH was found to increase mean number of shoots regenerated perexplant in case of internodal explants. Maximum number of shoots (6.33-7.5) were induced on MS2 medium with pH adjusted in the range of 5.8-6.8 (Figure 3B). But in case of leaf explants, maximum of four shoots regenerated per explants on medium with pH adjusted in range of 5.3-5.8 (Figure 3B).

Medium pH was also found to effect the chlorophyll content. It was found to decrease with increasing medium pH (Figure 3C-D). However, this change was significant in case of leaf explants. Maximum chlorophyll content was estimated in the leaf explants cultured on MS2 medium with pH adjusted in range of 5.3 to 5.8. It was noteworthy that chlorophyll a/b ratio in leaf explants remain uniform within the pH range of 4.8 to 6.8 (Figure 3D).

### Effect of silver nitrate

The effect of silver nitrate evaluated by culturing leaf and internodal explants of potato cv. ‘Kufri Chipsona 1’ on MS medium containing BA; 10 µM, GA_3_; 15 µM and different AgNO_3_ concentrations (5-20 µM) showed that explants responded differentially with varying concentrations of AgNO_3_ (Figure 1 and 4). The regeneration frequency from both the explants was observed to increase by about two folds with increase in AgNO_3_ concentration from 5 µM to 10 µM (Figure 4A). Maximum of 32.11% leaf explants and 59.99% internodal segments of ‘Kufri Chipsona1’ showed shoot organogenesis on MS1 medium containing BA (10 µM) and GA_3_ (15 µM) (here onwards referred to as MS2 medium). Any further increase in AgNO_3_ concentration lead to decreased shoot organogenesis in both explants of potato.

**Figure 4:**
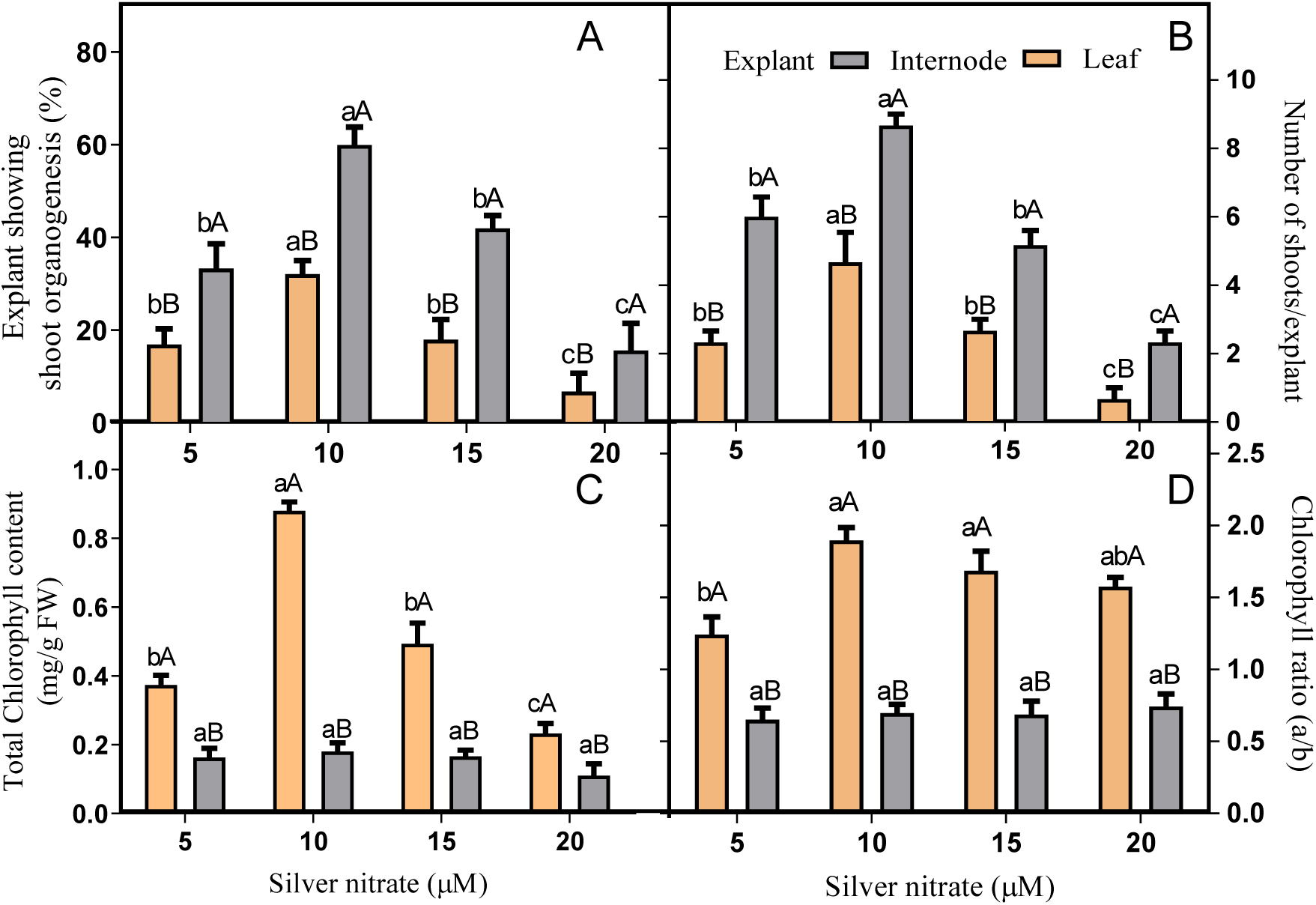
The effect of silver nitrate (AgNO_3_) on shoot organogenic efficiencies and associated chlorophyll changes in leaf and internodal explants taken from potato cultivar ‘Kufri Chipsona 1’. Data were recorded after 6 weeks and analysed by ANOVA. Mean values within the explants were compared using Duncan’s multiple range test DMRT and between the explants were compared using unpaired t-test at P< 0.05. Bars with same lowercase letter (within explants) and uppercase letter (between explants) were non-significant at P<0.05.

Response of both explants also varied from each other with respect to mean number of shoots per explant. The number of shoots regenerated from internodal explants was almost twice than shoots regenerated from leaf explants (Figure 4B). Maximum (8.67) shoots per explant were observed from internodal explants cultured on MS2 medium. Like regeneration frequency, mean number of shoots per explants was also found to decrease with further increase in AgNO_3_ concentration (Figure 4B).

Chlorophyll content and chlorophyll a/b ratio was also found to vary with concentration of silver nitrate in the medium (Figure 4C-D). Higher concentration of AgNO_3_ (above 15 µM) in the regeneration medium was observed to decrease total chlorophyll content (Figure 4C). However, the variation in chlorophyll a/b ratio with increase in AgNO_3_ ratio was not found to be significant for leaf explants cultured on regeneration medium containing AgNO_3_ above 10 µM (Figure 4D). The chlorophyll content or chlorophyll a/b in the internodes did not change with increase or decrease in AgNO_3_ concentration.

### Clonal fidelity studies

In the present study, indirect shoot organogenesis was achieved. Therefore, it was important to establish the clonal fidelity of the regenerated shoots vis-à-vis mother cultures. A total of 10 primers (5 RAPD and 5 ISSR) were tested and all the markers (both RAPD and ISSR) scored were found to be monomorphic (Figure 1C-D). A maximum of 5 bands (for RAPD primer 4) and minimum of 1 band (for many primers) were scored (Kaur et al. 2017) and size of amplified fragments ranged from 200 bp to 2000 bp. A total of 43 markers were scored (24 for RAPD and 19 for ISSR). All the regenerated plants were found to be true-to-type to that of mother cultures

## Discussion

Shoot organogenesis protocol, a pre-requisite for genetic manipulation of plants has been well documented in potato but is complicated by use of two step procedure (Yee et al. 2001, Yasmin et al. 2003, Elaleem et al. 2009, Molla et al. 2011, Rezende et al. 2013, Ghosh et al. 2014, Abdelaleem 2015, Campos et al. 2016). Moreover, genotype specific response of potato cultivars to plant growth regulators (PGRs) has also been reported by many researchers (Khokan et al. 2009, Pal et al. 2011, Wheeler et al. 1985). In this context, previously we have developed one step, genotype independent shoot organogenesis protocol for potato (Kaur et al. 2017). As a next logical step, the present study was focussed to optimise other factors to further improve the efficiency of the established protocol.

Shoot organogenesis is known to be dependent upon many factors such as medium composition and culture conditions (Kumar et al. 2003, Parris et al. 2012, Halloran and Adelberg 2011). In the present study, effect of medium composition and pH on shoot organogenesis was evaluated. The study was carried out to achieve the goal of maximum shoot regeneration with minimum efforts. Although the effect of various factors such as pH, photoperiod, silver nitrate, sucrose concentration, node length, media consistency and gelling agent on micropropagation and microtuberization has already been reported for various cultivars of potato (Sandra and Maira 2013, Sharma et al. 2011, Turhan 2004, Zobayed et al. 2001, Veramendi et al. 1997) and other plants (Fujiwara et al. 1995, Kaczperski et al. 1991, Erwin et al. 1991, Kozai and Iwanami 1988, Niu and Kozai 1997, Kozai et al. 1995, Adelberg and Toler 2004) yet no scientific evidence is available describing the effect of above mentioned factors on shoot organogenesis in potato cultivars. The relation of shoot organogenic frequencies and culture conditions reported here may provide experimental basis for development of efficient regeneration protocol in potato.

Medium pH and solidifying agent were found to play a vital role in deciding regeneration efficiencies in potato. Medium solidified using clarigel as gelling agent (Figure 1 and 2) and adjusted to pH in range of 5.3-5.8 (Figure 3) was found to be the most suitable for shoot organogenesis. In general, both leaf and internodal segments has potential to regenerate into whole plant but our study reveals that internodes are better explants and were found to be more tolerant towards changing culture conditions in comparison with leaf explants. A variation in regeneration potential of explants has been reported earlier in potato by some workers (Sabeti et al. 2013, Seabrook Douglass 2001, Yee et al. 2001, Sharma et al. 2014, Kumar et al. 2017). The present findings are in line with the previous reports describing better response from internodal explants over leaf explants for regeneration of plants (Sabeti et al. 2013). However, there are also reports highlighting better response from leaf explants as compared to internodal explants (Ali et al. 2007, Shirin et al. 2007). The explant specific differences could be due to the variations in the endogenous levels of plant growth regulators in these explants, which could have marked effect on morphogenesis (Krikorian 2000, Pal et al. 2011).

Generally, clarigel is used in much lower concentration as compared to agar or gelrite which provide more hydrating and nutritious environment to the explants (Kumar et al. 2003). This may be the reason for accerlated shoot regeneration and shoot length on medium solidified with clarigel. Furthermore, it has been reported that clarigel is free from phenolic compounds and impurities in comparison to other gelling agents (Scherer et al. 1988) and thus beneficial for growth and organogenesis (Kumar et al. 2003).

Ethylene accumulation in culture bottles under in vitro propagation systems seems to be a major problem in potato regeneration studies. It has been reported that potato is much sensitive to ethylene accumulation (Jackson et al. 1987). Therefore, a potent ethylene inhibitor (silver nitrate; AgNO_3_), reported to reduce hyperhydricity in sunflower (Mayor et al. 2003) and branching in potato (Turhan 2004) was used in regeneration studies. In the present study, the incorporation of AgNO_3_ was found to be mandatory for induction of shoot regeneration. It was observed that addition of AgNO_3_ (10 µM) in medium supplemented with optimum concentration of BA and GA_3_ lead to maximum shoot organogenesis in CS-1 (Figure 4). The use of AgNO_3_ to increase regeneration frequencies have been reported in other plants such as cassava, pomegranate, alfalfa etc. (Zhang et al. 2001, Naik and Chand 2003, Li et al. 2009).

Chlorophyll content is used as an indicator of plant health (Sarropoulou et al. 2016). In the present study, maximum chlorophyll content and ratio (a/b) was observed from leaf explants cultured on medium containing AgNO_3_ (10 µM), BA (10 µM) and GA_3_ (15 µM) adjusted to pH range of 5.3-5.8. Increase or decrease in AgNO_3_ concentration or medium pH lead to reduction in chlorophyll levels (Figure 4). It has been previously reported that presence of any abiotic stress results in chlorophyll degradation which ultimately reduce chlorophyll content in the explants (Chen and Djuric 2001). It was noteworthy that gelling agent was not having much effect on chlorophyll content of the explants.

Since shoot organogenesis was achieved through intermediate callus phase, therefore, clonal fidelity using regenerated shoots was tested using RAPD and ISSR markers. The utility of molecular markers in studying clonal fidelity of in vitro propagated plants is well documented (Khatab and El-Banna 2011, Kumar et al. 2010, Rani and Raina 2000, Nag et al. 2012, Mehta et al. 2011). All the scored markers (both by RAPD and ISSR) were found to be monomorphic indicating the genetic uniformity of regenerated plants (Figure 1C-D). Probably this could be due to short exposures time in the culture medium. Earlier, the length of culture (Bairu et al. 2006, Rodrigues et al. 1998) and the presence of a disorganized growth (callus) phase in tissue culture were considered as major factor that may cause somaclonal variation (Rani and Raina 2000).

In conclusion, an efficient shoot organogenesis protocol was developed for potato cultivar CS-1 highlighting the importance of medium composition and culture conditions. The novelty of developed protocol have potential to significantly influence crop improvement programmes in potato.

## Acknowledgement

Amanpreet Kaur is thankful to University Grants Commission (UGC), New Delhi for the award of Maulana Azad National Fellowship for minority students. TIFAC-CORE is thanked for the facilities to carry out research work.

## Author contribution

Amanpreet Kaur conducted the experiments, compiled data, analyzed the results and wrote the initial draft of the manuscript, Amanpreet Kaur and Anil Kumar conceived and designed the experiments, Anil Kumar finalized the manuscript.

